# ExplorePipolin: reconstruction and annotation of bacterial mobile elements from draft genomes

**DOI:** 10.1101/2022.06.18.496689

**Authors:** L. Chuprikova, V. Mateo-Cáceres, M. de Toro, M. Redrejo-Rodríguez

## Abstract

**Motivation:** Detailed and accurate analysis of mobile genetic elements (MGEs) in bacteria is essential to deal with the current threat of multiresistant microbes. The overwhelming use of draft, contig-based genomes hinder the delineation of the genetic structure of these plastic and variable genomic stretches, as in the case of pipolins, a superfamily of MGEs that spans diverse integrative and plasmidic elements, characterized by the presence of a primer-independent DNA polymerase.

**Results:** ExplorePipolin is a Python-based pipeline that screens for the presence of the element and performs its reconstruction and annotation. The pipeline can be used on virtually any genome from diverse organisms and of diverse quality, obtaining the highest-scored possible structure, and reconstructed out of different contigs if necessary. Then, predicted pipolin boundaries and pipolin encoded genes are subsequently annotated using a custom database, returning the standard file formats suitable for comparative genomics of this mobile element.

**Availability:** All code is available and can be accessed here: github.com/pipolinlab/ExplorePipolin

## 1. Introduction

Infectious diseases caused by multidrug-resistant pathogens pose a major threat to global health, food security and development today, a situation aggravated by the insufficient production of new drugs (Zhu *et al*., 2021; Murray *et al*., 2022). Antimicrobial resistance (AMR) and virulence factors are largely encoded by itinerant genome stretches known as mobile genetic elements (MGEs) that act as a platform for their transference and spreading (Partridge *et al*., 2018; von Wintersdorff *et al*., 2016). Thus, MGEs constitute an essential part of bacterial genome and can contribute to their evolution, fitness, and pathogenicity. However, despite recent efforts, many MGEs are overlooked or misannotated in population genomics analyses and they are generally underestimated (Oliveira Alvarenga *et al*., 2018; Hua *et al*., 2021; Ross *et al*., 2021).

The recent accessibility of high-throughput sequencing methods as part of surveillance programs of bacterial pathogens allows the genomic and metagenomic monitoring of the expansion of bacterial strain-specific markers, including virulence and AMR genes. However, these fast-evolving methods also generated a burden of data that must be throughout processed and analyzed (Mitchell and Simner, 2019). This is particularly problematic regarding the study of dynamics and plasticity of MGEs, as they can range in size from very simple and small elements, such as insertion elements (IS), coding for only the transposase necessary for their relocation, to large prophages, transposons and plasmids, which can be tens or hundreds of kilobase pairs in length and also appear associated among themselves (Partridge *et al*., 2018; Durrant *et al*., 2020; Benler *et al*.). Further, MGEs prediction and analysis is hindered by their great modularity and rapid evolution through gene acquisition and gene loss. Many pipelines designed for the analysis of MGEs are specialized and rely on the identification of hallmark genes, like plasmid replication proteins (Carattoli and Hasman, 2020), relaxases (Alvarado *et al*., 2012) or specific transposase or recombinases for integrative elements (Ross *et al*., 2021; Siguier *et al*., 2012; Moura *et al*., 2009; Cury *et al*., 2016, 2017). Some works have focused on the use of high-quality reference genomes, but at the cost of diversity loss (Jiang *et al*., 2019). Similarly, a few recent methods for annotation of prophages are able to identify insertion boundaries on complete genomes (Guo *et al*., 2021; Arndt *et al*., 2019). However, the great majority of currently available genomes are based in short sequencing reads technologies, resulting in draft genomes, made up of tens or hundreds of heterogeneous contigs. This makes accurate tracking of the integrated elements virtually impossible (Arredondo-Alonso *et al*., 2017). The usage of raw reads to scaffold integrative elements has been successfully applied to reconstruct some elements (Durrant *et al*., 2020), though it entails large computational resources, hindering its application for massive screenings. More recently, the conserved sequences of the transposase binding sites and their unique architecture are shown to carry a signal that is sufficient to delineate the 5′ and 3′ boundaries of Tn7-like elements (Benler *et al*.). Pipolins are a novel group of MGEs found in diverse major bacterial phyla but also mitochondria (Redrejo-Rodríguez *et al*., 2017). The only gene shared by all pipolins encodes for a subgroup of DNA polymerases from family B, named piPolB (for *p*rimer-*i*ndependent *PolB*). This relates pipolins to other elements commonly referred as “self-synthesizing” by the presence of a PolB, the eukaryotic polintons and casposons, detected in archaea and some bacteria (Krupovic *et al*., 2014; Kapitonov and Jurka, 2006; Krupovic and Koonin, 2016).

Pipolins in *E. coli* are diverse and plastic but, besides the piPolB gene, they share conserved att-like direct repeats (hereafter *atts*), overlapping with a tRNA gene. We previously took advantage of that common structural features to delineate their structure out of contig-scale *E. coli* draft genomes (Flament-Simon *et al*., 2020) using a preliminary customized Python pipeline (Chuprikova, 2020). However, beyond *E. coli*, the *att* sequences are not conserved and indeed, pipolins have been identified as circular plasmids without direct repeats in several Fungi mitochondria as well as in a number of bacteria, spanning very diverse species, like *Enterobacter hormaechei, Staphylococcus epidermidis* or *Lactobacillus fermentum* (Redrejo-Rodríguez *et al*., 2017).

In this work, we now present ExplorePipolin, a full-fledge Python-based pipeline, that allows robust reconstruction and homogeneous characterization of pipolins from diverse bacterial genomes. The pipeline analyzes each contig for the presence of a pipolin marker and proposes the highest-scored possible structure, reconstructed out of different contigs if necessary. Then, the element is delimited by predicted *atts* and pipolin encoded genes are subsequently annotated using a custom database. The workflow is publicly available and can be installed from its repository on the GitHub or via Conda. It aims to be a documented, composable, and reusable application (Brack *et al*., 2022), easily translatable to other elements, such as ICEs or casposons, among others.

## 2. Methods

### 2.1 Pipeline outline

As pipolins are defined by the presence of a piPolB gene (Redrejo-Rodríguez *et al*., 2017), ExplorePipolin pipeline workflow (Figure 1) must start by searching the (multi)fasta query sequence for a piPolB coding sequence. To achieve high sensitivity and accuracy in this limiting step, we updated the previous dataset of diverse piPolBs (Redrejo-Rodríguez *et al*., 2017) using Jackhmmer searches on several databases (Potter *et al*., 2018) and generated an HMM profile spanning all possible diversity. This profile was then used to scan for piPolBs in the translated fasta nucleotide sequences using as cutoff an E-value of 10^−50^, selected to detect all known piPolBs and reduce the false positives hits from sequences that belong to pPolBs and other related B-family DNA polymerase groups (Kazlauskas *et al*., 2020).

**Figure 1.**
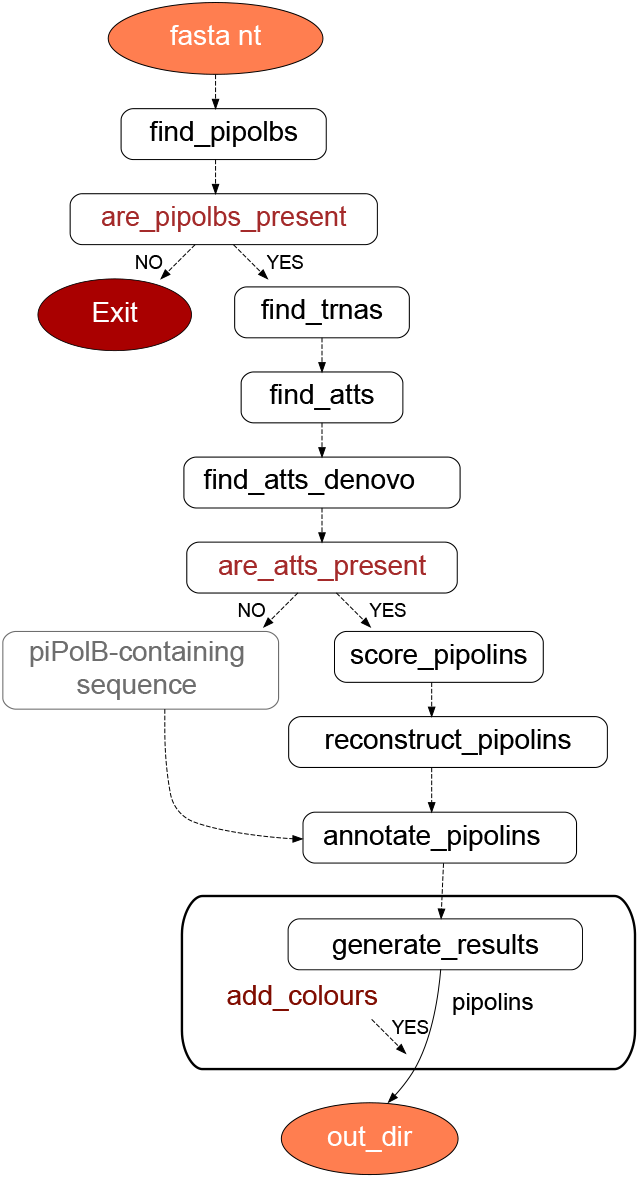
Workflow of ExplorePipolin. Main nodes and tasks are indicated. Dataflow throughout the pipeline is managed with Prefect (https://www.prefect.io/).

When a piPolB-containing fragment is identified, DNA sequences are subsequently scanned for the pipolin boundaries. First, as pipolins often integrate on tRNA genes, we search for nearby tRNA or tmRNA gene, using the Aragorn tool (Laslett and Canback, 2004) and that will be as a verification step for the identification of the *att*-like direct repeats. In *E. coli*, pipolin *atts* are highly conserved among diverse strains (Flament-Simon *et al*., 2020), which allow their quick identification with BLAST using the sequence of the *attR* from *E. coli* strain 3-373-03_S1_C2 pipolin (Redrejo-Rodríguez *et al*., 2017; Richter *et al*., 2018). Alternatively, when known *att* sequences are not found, we search for “de novo” direct repeats that might participate in the pipolin mobilization. Briefly, we first look for short direct repeats around a piPolB gene using BLAST (85% identity with word size 6, plus strand only to exclude inverted repeats). This strategy is also implemented in other tools, as Phaster (Arndt *et al*., 2019). All the predicted repeats are annotated and situations with an unexpected number or repeats (i.e., one or more than two) are handled during the element reconstruction (see below). To facilitate the comparison among diverse pipolin genomic structures, we stablished that whenever an *att* overlapped with a tRNA, that will be the *attR* and the other repetition will be denoted as *attL*. Importantly, unsuccessful repeats identification will not lead to the pipeline failure. Notwithstanding technical issues, as mentioned above, pipolins have been also identified as circular plasmids. Thus, in the absence of known or predicted repetitions, the boundaries of the pipolin are arbitrarily settled in a maximum of 30 kb at each side of the piPolB gene (Figure 2C, bottom pipolin), which may be modified by the user (option --max-inflate), to allow a downstream study of a sequence that might constitute either a plasmidic pipolin or a pipolin-containing fragment.

**Figure 2.**
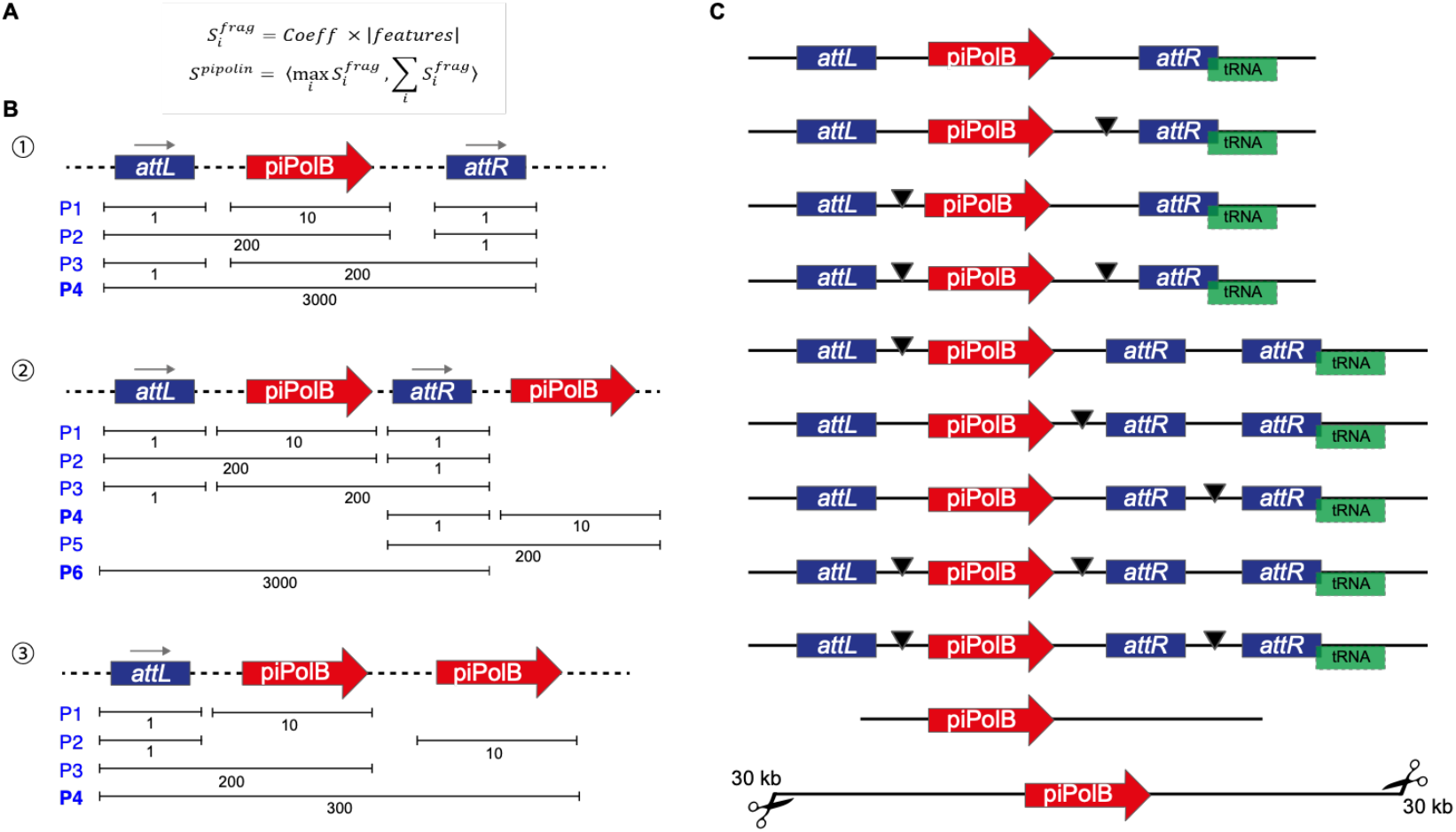
Featured-based mobile element reconstruction from draft genomes. **A.** Description of the scoring applied to possible pipolin recontructions. First, (i) the score of each fragment 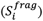 is calculated as a coefficient multiplied by the number of features. The coefficient depends on the feature present in the fragment: 1000 for piPolB surrounded by atts, 100 for piPolB with *atts* from one side, 10 when piPolB only is present, 1 for *atts* only. Then, (ii) the score of each alternative predicted pipolin (*S*^*pipoline*^) is a tuple (i.e., an ordered pair) of two numbers: 1) the maximal score among all fragments and 2) the sum of the scores of all fragments. **B**.Scoring of alternative pipolin reconstructions. Features are linked with a dashed line as reconstruction may use features from one or multiple contigs. The maximal score of the generated pipolin alternatives is indicated. In the simplest example (1), when the piPolB and two att sequences can be identified on the same contig, reconstruction will lead to the P4 alternative. In case of fragmented pipolin, recostruction will lead to P3, P2 or P1 alternative respective to the location of disruptions. As indicated with rightwards arrows, the orientation of the piPolB and the categorization of atts as attL and attR is established as detailed in the text. In more complex scenario (2), several pipolins may be possible. In this case, P6 has the highest score (3000, 3000). After choosing P6 and removing overlapping variants, P4 would be included in the output as a second pipolin, containing only piPolB. In the case (3), four pipolins are possible, but the P4 reaches the highest score by the sum of the features included into the fragment. **C**.Examples of most common reconstructed output pipolins, containing or not the *atts* and the overlapping tRNAs. The black arrowhead denotes an assembly gap introduced after the reconstruction step in order to create a single pipolin out of individual fragments located on different contigs.

### 2.2 Element reconstruction procedure

As mentioned above, the generalized use of short-reads high-throughput sequencing methods, contig-based bacterial genomes or assemblies are still the most common bacterial genome format nowadays. These fragmented genomic sequences pose a great difficulty to study the structure and plasticity of MGEs of middle-large size as pipolins (10-40 kb), as the element can be often split into two or more contigs. Scaffold of pipolins might be improved using raw reads or comparison with related elements, as they maintain a certain degree of gene synteny (Redrejo-Rodríguez *et al*., 2017; Flament-Simon *et al*., 2020). However, those approaches entail some risks and require high computational resources, thus hindering quick analysis of new draft genomes and/or high throughput screenings and tend to perpetuate previous mistakes. We decided to rely on the information available in the genome draft to reconstruct the possible pipolin structures and score them based on genetic and bioinformatics criteria to efficiently obtain the most likely pipolin structure. We only granted two assumptions; first that two pipolins cannot be overlapping and, second, piPolB gene is the center of the pipolin, thus there is no pipolin only with *atts*, but there can be a pipolin without an *att*, being *att-piPolB-att* the highest scored structure. Using these criteria (Figure 2), the pipeline iteratively leaves pipolins with the best score and removes those overlapping them. With this approach, ExplorePipolin could be used for a wide diversity of pipolin structures, from the already studied ones in *E. coli* (Flament-Simon *et al*., 2020) to a broad range present in other bacterial genomes (see below).

## 3. Results

### 3.1 Pipeline implementation

ExplorePipolin is available on GitHub under GPL-3.0 license. It can be installed from source files and as a Conda package. It is a simple application that only requires a minimal familiarity with Linux-based commandline environments.

After reconstruction, the final most likely pipolin will be selected and the discarded structures are removed by default, although they can be optionally kept for further exam (option --keep-tmp). Additionally, when alternative orientations of the piPolB-containing fragment are possible, both reconstructions are generated and denoted as v0 for that with the piPolB gene pointing toward the *attR* (Figure 2), which is the most common situation in *E. coli* pipolins (Flament-Simon *et al*., 2020), and alternative pipolins will be referred as v1, v2…

The pipolin reconstructed structure(s) are subsequently annotated using a custom Prokka-based pipeline that includes previously known pipolin-related genes. The output includes files in standard formats GBK and GFF generated for wide compatibility for downstream analysis of pipolins genetic structure. An additional file, appended with the suffix “single_record”, contains a final reconstructed pipolin comprising individual pipolin fragments from different contigs joined by an assembly gap feature type. The annotated *att* direct repeats are also included in the output files and labeled as “pipolin conserved” or “de novo”, according to their relationship to known *att*-like sequences of *E. coli* pipolins or identified as a novel direct repeat fragment.

Moreover, we also include color information of common pipolin features (i.e. piPolB, *att*, tyrosine recombinases, restriction-modification systems, uracil DNA glycosylase…) for straightforward representation with EasyFig (Sullivan *et al*., 2011). Default colors from previously used scheme can be modified providing a TSV file (option --colours) or removed (--skip-colours). We found EasyFig a simple customizable visualization alternative, suitable for comparison of genetic structure and diversity of pipolins and convenient for non-bioinformaticians. We have also tested other recent related applications such as the web-based clinker (Gilchrist and Chooi, 2021) or the R package gggenomes (Hackl and Ankenbrand). The latter seemed enough versatile for pipolins representation (Figure S1). We have included in the GitHub repository scripts for plotting ExplorePipolin output in R using the gggenomes. In short, a custom Python script adapts the GBK files with some required modifications and calculates pipolin synteny and GC content according to the gggenomes recipe. Then, an R script uses these data to plot the sequences and the desired features. It is only necessary the final pipolin GBK files (“single_record”), a file specifying the order of the pipolins to compare in the representation, and the software requirements specified in the README file.

### 3.2 Examples of use

We have successfully applied the pipeline to diverse genomic sequences. Complete analysis of a medium size bacterial draft assemblies (5-6 MB) takes approximately 30-60 seconds per genome using multiple threads for annotation (option --cpus). Further optimization may be achieved using additional parallelization strategies, like GNU Parallel (Tange, O.). Moreover, most time-consuming step is the annotation of the reconstructed elements, and it can be passed up (option --skip-annotation) for saving time, for example, when large screening of thousands of genomes is done.

Notwithstanding a throughout modification of the original pipeline, ExplorePipolin analyzed the genomes of our laboratory *E. coli*-harboring pipolins collection obtaining nearly identical pipolin structures than in the original analysis (Flament-Simon *et al*., 2020) (Figure S1).

Furthermore, the new reconstruction strategy allows successful reconstruction and analysis of pipolins from a wide range of distant organisms, spanning different bacteria phyla and Fungi genomes (Figure 3). The usage of HMM profile searches gives rise to a highly sensitive and specific identification of piPolB coding sequences that are subsequently delineated and annotated, regardless of the presence of the known *att* sequences or any direct repeat at all.

**Figure 3.**
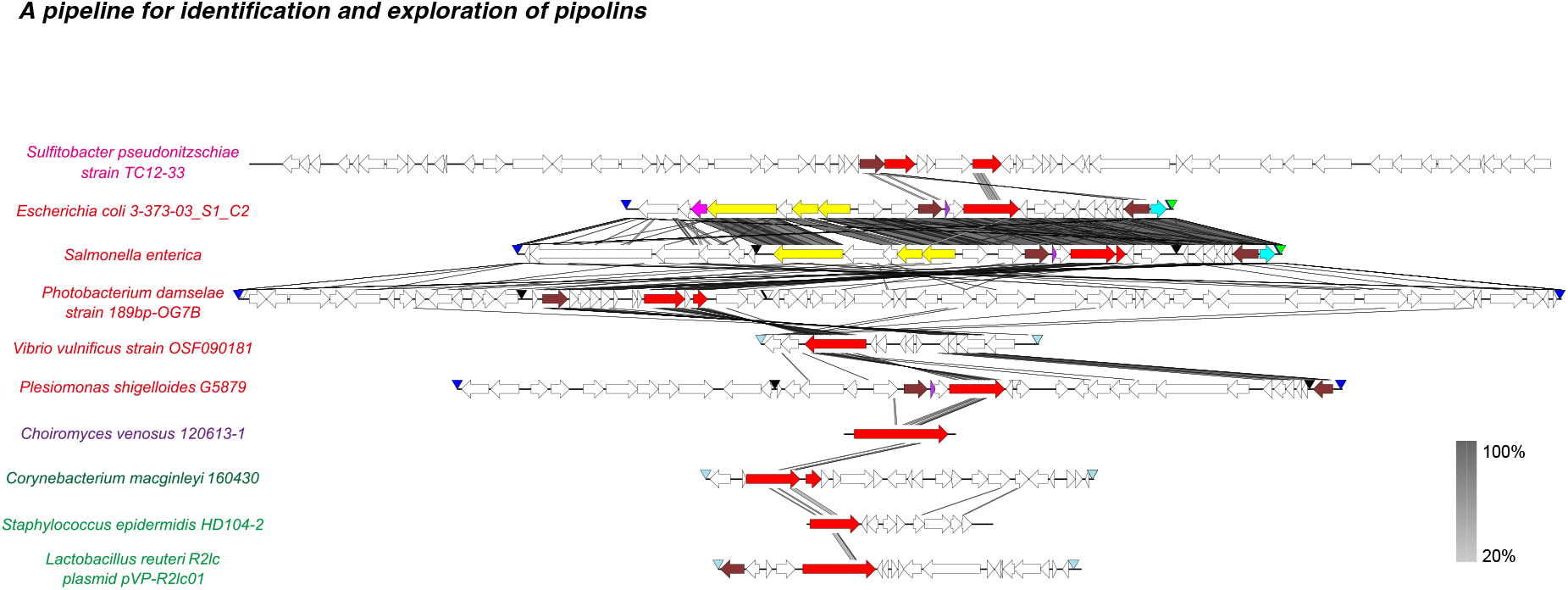
Genetic structure of diverse pipolins from genomes from a wide range of bacteria reconstructed by ExplorePipolin. Predicted protein-coding genes are represented by arrows, indicating the direction of transcription and more common genes are colored following Prokka annotation (piPolB in red, tyrosine recombinase in brown, UDG in cyan, excisionase in purple and metallohydrolase in magenta). When detected, other features are indicated as colored arrowheads: sequence gaps (black), *E. coli* related-*atts* (navy blue), *de novo* detected direct repeats (steel) and tRNAs (green). The greyscale on the right reflects the percent of amino acid identity between pairs of sequences. The image was generated by EasyFig software using tBlastX for elements comparison. Names of the analized genomes are indicated on the left and colored by taxonomy: magenta, Alphaproteobacteria; red, Gammaproteobacteria; purple, fungi, forest green, Actinobacteria and green, Firmicutes.

## 4. Conclusion

ExplorePipolin is an example of a comprehensive pipeline that not only looks for MGE presence by marker screening but also reconstructs the whole pipolin element structure and attempt to provide precise boundaries, making further analysis more accurate and straightforward. Importantly, it can be widely used to screen for pipolins on virtually any genome of a wide range of organisms and diverse assembly qualities and it returns putative reconstructed pipolins with homogenous annotation suitable for comparative genomics of this mobile element.

Furthermore, the modular structure of the pipeline will facilitate its future update for bulk analysis of other elements with one or few universal hallmark features, particularly those related with pipolins, such as casposons or polintons.

## Acknowledgements

We thank Mario R. Mestre for the generation of the piPolBs HMM profile. We are also grateful to members of the MR-R lab for discussions and suggestions.

## Funding

This work has been supported by the Spanish Ministry of Science, Innovation and Universities, Grant Number PGC2018-093723-A-I00 (AEI and FEDER, UE) to M.R.R.

### Conflict of Interest

none declared.

---

**Supplemenary Figure 1.**
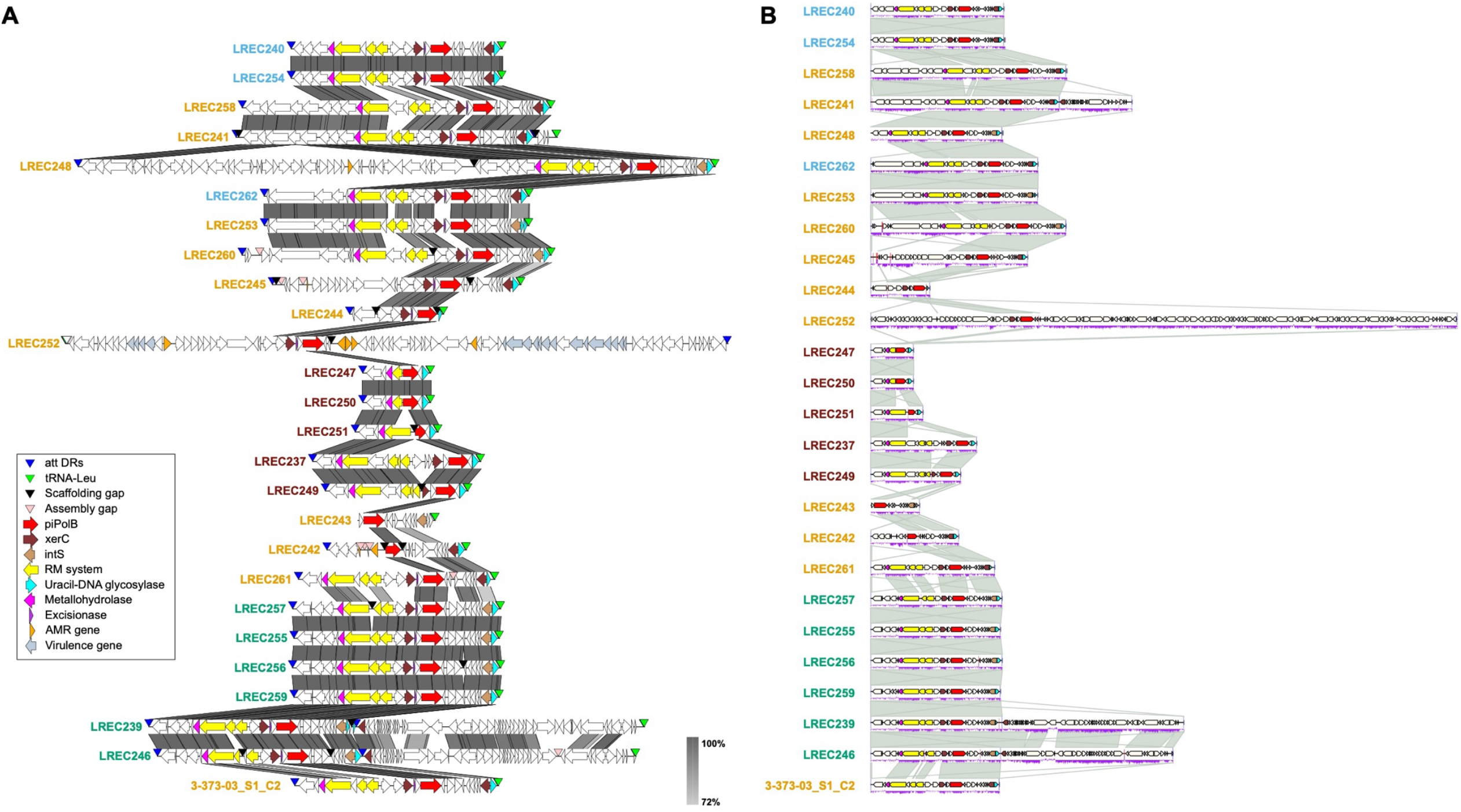
Comparison between *E. coli* pipolin structures from LREC collection (Flament-Simon et al. 2020) as originally delineated (A) and with the current version of ExplorePipolin (B). Pipolins in panel B were plotted using gggenomes (Hackl,T. and Ankenbrand,M.).

## Notes

### Competing Interest Statement

The authors have declared no competing interest.

