## Supplementary figures and images for "ExplorePipolin: reconstruction and annotation of bacterial mobile elements from draft genomes"

### Figure S1

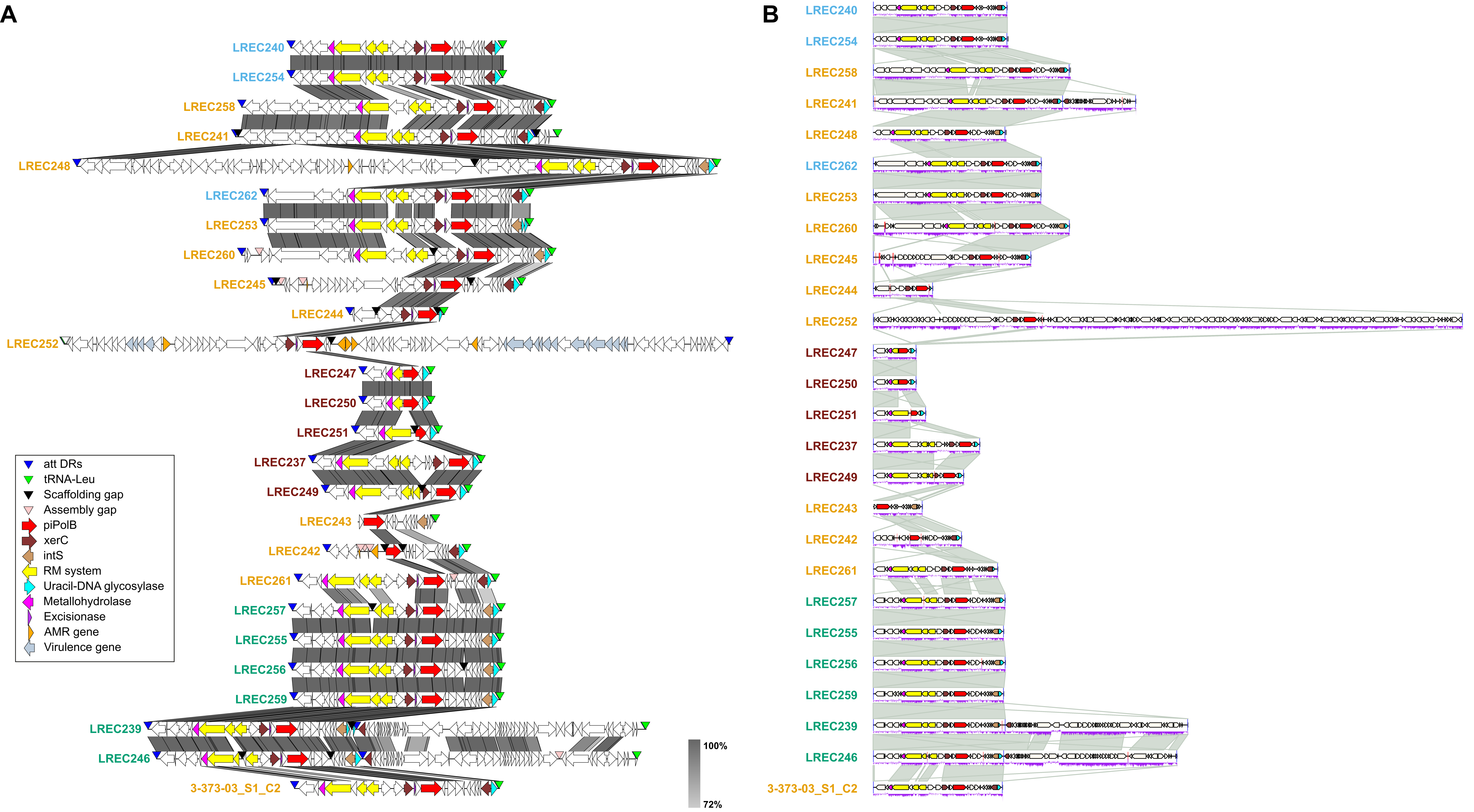
